# A core signaling mechanism at the origin of animal nociception

**DOI:** 10.1101/185405

**Authors:** Oscar M. Arenas, Emanuela E. Zaharieva, Alessia Para, Constanza Vásquez-Doorman, Christian P. Petersen, Marco Gallio

## Abstract

All animals must detect noxious stimuli to initiate protective behavior, but the evolutionary origin of nociceptive systems is not well understood. Here, we show that a remarkably conserved signaling mechanism mediates the detection of noxious stimuli in animals as diverse as flatworms and humans. Planarian flatworms are amongst the simplest bilateral animals with a centralized nervous system, and capable of directed behavior. We demonstrate that noxious heat and irritant chemicals elicit robust escape behaviors in the planarian *Schmidtea mediterranea*, and that the conserved ion channel TRPA1 is required for these responses. TRPA1 mutant fruit flies (*Drosophila*) are also defective in the avoidance of noxious heat ^1-3^. Unexpectedly, we find that either the planarian or the human TRPA1 can restore noxious heat avoidance to TRPA1 mutant *Drosophila*, even though neither is directly activated by heat. Instead, our data suggest that TRPA1 activation is mediated by H_2_O_2_/Reactive Oxygen Species, early markers of tissue damage rapidly produced as a result of heat exposure. Together, our data reveal a core function for TRPA1 in noxious heat transduction, demonstrate its conservation from planarians to humans, and imply that human nociceptive systems may share a common ancestry with those of most extant animals, tracing back their origin to a progenitor that lived more than 500 million years ago.

---

Each animal group uses specialized sensory systems to detect and avoid predators, find food sources and mates. Due to specific demands, sensory systems evolve independently in different species. Yet, while each animal group lives in a sensory world that is essentially unique, most of what we know about sensory representation comes from a very limited number of species - a handful of vertebrate and invertebrate model systems.

The detection of potentially harmful conditions is a core sensory task. The ion channel TRPA1 is remarkably conserved across animal evolution and has been implicated in the response to a broad range of electrophilic irritant chemicals as well as to noxious hot or cold temperature in humans ^4^, mice ^5-11^, and flies ^2,3,12^. Interestingly, while TRPA1’s sensitivity to irritant chemicals has been widely conserved (^13^, in all but the *C. elegans* homolog ^14^), its temperature gating appears to have changed repeatedly during evolution ^15^. *In vitro*, some mammalian TRPA1 homologs are activated by noxious cold ^16^, while others are insensitive to temperature (reviewed in ^15^). In contrast, TRPA1 from chicken ^17^, various reptiles, and *Xenopus* frogs are activated by warm temperatures ^18,19^, and the zebrafish genome encodes two distinct paralogs, only one of which shows thermosensory responses ^20,21^. The situation in invertebrates is also complex: while *C. elegans* TRPA1 is activated by cold ^14^, insect TRPA1s (honeybee ^22^, mosquito ^23^ etc.) are activated by warm temperatures, and the fruit fly homolog is spliced into at least four variants ^3^, including both heat-sensitive and heat-insensitive ones ^3,12,24^.

What is the ancestral function of TRPA1? How ancient is its association with nociceptors? Planarian flatworms are an attractive system in which to study evolutionary origins of sensory transduction. As members of the phylum Platyhelminths, planarians are considered among the simplest animals with bilateral symmetry and a centralized nervous system. From an evolutionary perspective, they are nearly equally distant from taxa that include species extensively studied such as nematodes, flies and mice (^25^ and Figure 1a). Furthermore, recent work on regeneration has led to the development of RNAi protocols to systemically knock-down the expression of selected genes *in vivo* ^26^. Here, we use these tools to investigate the function of TRPA1 in the freshwater planarian *Schmidtea mediterranea*.

**Figure 1:**
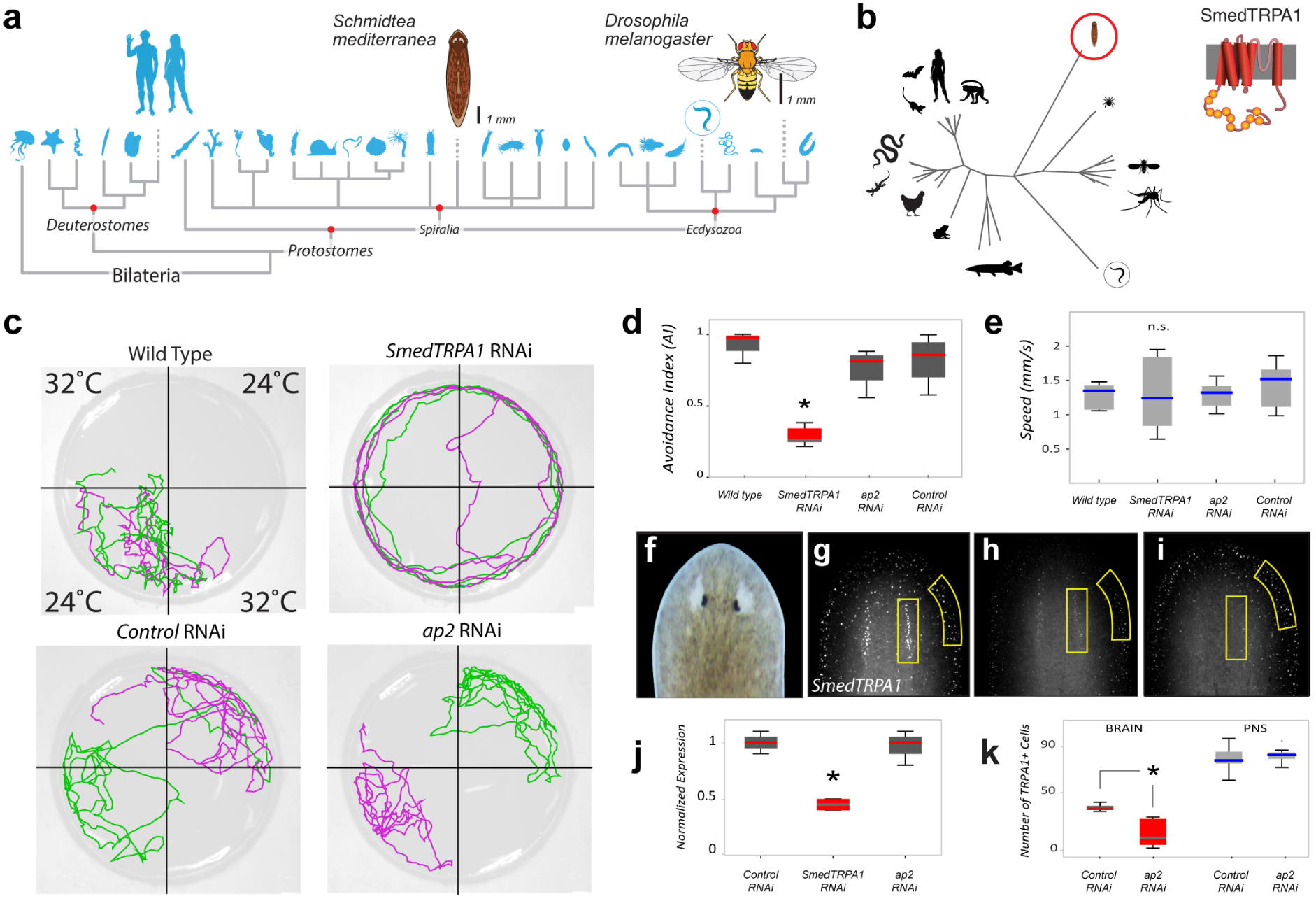
*Smed-TRPA1* is required for noxious heat avoidance in the planarian worm *S. mediterranea.* **a**) Phylogeny of Bilateria, showing the position of *Schmidtea* (*C. elegans* is circled). **b**) Phylogenetic tree constructed from an alignment of full-length TRPA1 protein sequences from a variety of species, Smed-TRPA1 is circled and a model of the channel’s structure is shown (circles=ankyrin repeats, cylinders=transmembrane domains). **c**) 2-choice assay for heat avoidance. In each trial two opposing floor tiles are set to 24°C and two to 32°C (noxious heat). Tracks of two worms during one such trial are shown in green and purple. Unlike wild-type, controls (*unc22* RNAi), and *ap2* RNAi, *Smed-TRPA1* RNAi animals were not confined to the cool quadrants. **d**) Avoidance index for 32°C for RNAi animals. *Smed-TRPA1* RNAi animals show a reduced avoidance index for heat (N= 5 groups of 10 animals, *P< 0.05, ANOVA). **e**) *Smed-TRPA1* RNAi does not impact the animal’s speed of movement (N=10 animals; n.s. = not significantly different). **f-i**) *In situ* hybridization with a *Smed-TRPA1* probe in (**g**) Control (*unc22*) RNAi, (**h**) *Smed-TRPA1* RNAi and (**i**) *ap2* RNAi animals (head region, see **f**), demonstrates overall reduction of mRNA by *Smed-TRPA1* RNAi (independent quantification by Q-PCR is shown in **j;** N=4 replicates of 3 animals each, * = P<0.05, ANOVA). **k**) In contrast, *ap2* RNAi reduces the number of *Smed-TRPA1*-expressing cells in the brain region, but not in the periphery (N=9 animals, * = P< 0.001, t-test); in all box plots, edges of the boxes = first and third quartiles, line = median, whiskers = data range, crosses = outliers.

A fragment of the *S. mediterranea* TRPA1 gene has been previously used in *in situ* hybridization experiments as a marker for a subset of differentiated neurons ^27^. Starting from it, we cloned a full-length coding sequence for the gene (see methods for details), henceforth referred to as *Smed-TRPA1* (Figure 1b). To test whether *Smed-TRPA1* mediates the avoidance of noxious heat in *S. mediterranea*, we designed a two-choice avoidance assay (**Extended Data Figure 1**) based on the one we previously developed for fruit flies ^28^. Animals were introduced into a small circular chamber covered by a thin film of water, and tracked while making a choice between floor tiles kept at moderate (24°C) or hot (32°C) temperatures; the time spent in each quadrant was then quantified to calculate an avoidance index (AI). In this assay, *S. mediterranea* showed robust avoidance of heat (32°C, AI∼1), manifested as sharp turns away from the hot quadrants (Figure 1c). This is consistent with a nocifensive behavior, and indeed with the fact that *S. mediterranea* comes from cool water environments and can die from brief exposure to 35°C (^29,30^ and data not shown).

Remarkably, the avoidance of hot quadrants was severely disrupted by RNAi knock-down of *Smed-TRPA1* (Figure 1c-d and see **Extended Data Video 1**). *Smed-TRPA1* RNAi animals glided around the chamber without turning at the hot-cold boundaries (see tracks in Figure 1c), and ended up spending nearly equal time in hot as in cool quadrants. This is in sharp contrast with the behavior of both untreated and control worms (i.e. worms fed dsRNA targeted to a sequence not present in the worm genome, Figure 1c-d). Importantly, *Smed-TRPA1* RNAi worms glided around the chamber at comparable speed as controls (Figure 1e), and displayed robust negative phototaxis when given a choice between light and dark (in an independent assay, see **Extended Data Figure 2**), indicating that *Smed-TRPA1* RNAi does not mar gross locomotor functions, nor does it impact all aversive behavior.

**Figure 2:**
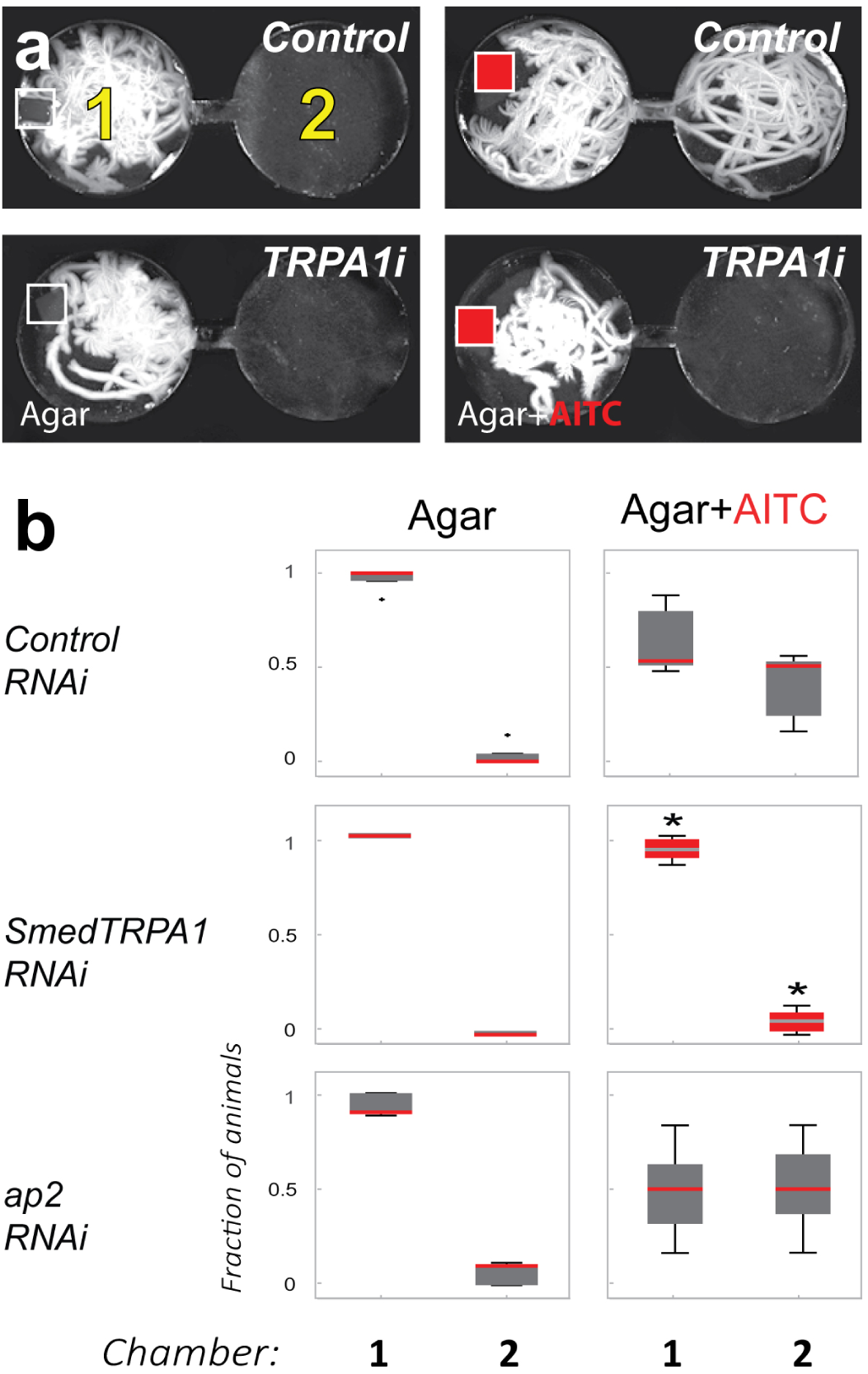
*Smed-TRPA1* is required for behavioral avoidance of the irritant chemical AITC. **a**) Two-chamber arena designed to quantify behavioral avoidance of chemical agonists of TRPA1. Planarian worms are introduced in chamber 1 in the presence of a mock Agar pellet (empty squares) or Agar+AITC (50 mM; red squares); their movement is then recorded for 5 minutes. The panels are maximum-projections of 5′ movies, illustrating the extent of worm movement (white tracks). **b**) In the presence of agar alone, control (*unc22*), *ap2* and *Smed-TRPA1* RNAi worms do not readily cross the narrow channel connecting chambers 1 and 2. In the presence of AITC, both control (*unc22*) and *ap2* RNAi worms exit chamber 1 and explore chamber 2. In contrast, *Smed-TRPA1* RNAi animals overwhelmingly remain in chamber 1 (N= 5 groups of 10 animals, fraction was calculated on the last 1′ of video; * = P<0.01, ANOVA comparing fraction of animals in chamber 1 or 2 across treatments; edges of the boxes = first and third quartiles, line = median, whiskers = data range, crosses = outliers).

RNAi knock-down of the transcription factor AP2 has been previously shown to impair the expression of TRPA1 in *S. mediterranea* ^27^. Based on this, we reasoned that *ap2* RNAi could provide an independent means to assess the role of TRPA1 in heat nociception. Unexpectedly, *ap2* RNAi animals did not display an avoidance defect, and instead avoided the hot quadrants as robustly as controls (Figure 1c-d). *In situ* hybridization revealed that *ap2* RNAi was effective in knocking-down *Smed-TRPA1* expression only within the brain, and not in peripheral neurons (Figure 1 f-k). The fact that animals with significantly reduced *Smed-TRPA1* expression within the brain behave normally suggests that *Smed-TRPA1* is required at the periphery for the detection or responses to noxious heat.

Next, we tested *Smed-TRPA1* knock-down animals for potential defects in chemical nociception, by assaying behavioral responses to Allyl isothiocyanate (AITC). AITC is the agent responsible for the pungent taste of mustard and wasabi, and a well-known chemical agonist of TRPA1 ^6,7,11^. We developed an arena consisting of circular chambers interconnected by small corridors that are not readily traversed by the worms (Figure 2a). Animals fed control dsRNA (see above) or *Smed-TRPA1* dsRNA were introduced in the first chamber in the presence of a mock agar pellet, or-alternatively-of an agar pellet laced with AITC; their behavior was then monitored for 5 minutes. Mock pellets were readily explored by untreated animals as well as by RNAi controls, which as a result remained in their vicinity. In contrast, AITC produced strong aversive responses including rapid withdrawal and abrupt turns. The worms ultimately escaped away from the chamber containing the pellet traversing the narrow corridors (Figure 2b and **Extended Data Video 2**). Again in sharp contrast to controls, *Smed-TRPA1* knock-down animals did not display aversive responses and instead remained in the first chamber, in the vicinity of the AITC-laced pellet (Figure 2b).

Our experiments show that *Smed-TRPA1* is a key mediator of both heat avoidance and chemical nociception *in vivo* in *S. mediterranea*. To begin studying the biophysical properties of the channel *in vitro*, we next performed whole cell patch-clamp electrophysiology on heterologously expressing cells. To achieve functional expression of Smed-TRPA1, we chose *Drosophila* S2 cells, a system previously used for *Drosophila* TRPA1 ^31^. Our recordings show that, in S2 cells, Smed-TRPA1 was activated by AITC (Figure 3). In contrast, the channel was not directly gated by heat (Figure 3a-d). Even when mis-expressed *in vivo*, in transgenic *Drosophila* (i.e. in all fly neurons), Smed-TRPA1 could be readily activated by AITC, but not heat (Figure 4). As a control, the *Drosophila* TRPA1-A variant ^3^ was activated by both AITC and heat in both contexts (Figure 3c-d and Figure 4, and ^32^).

**Figure 3:**
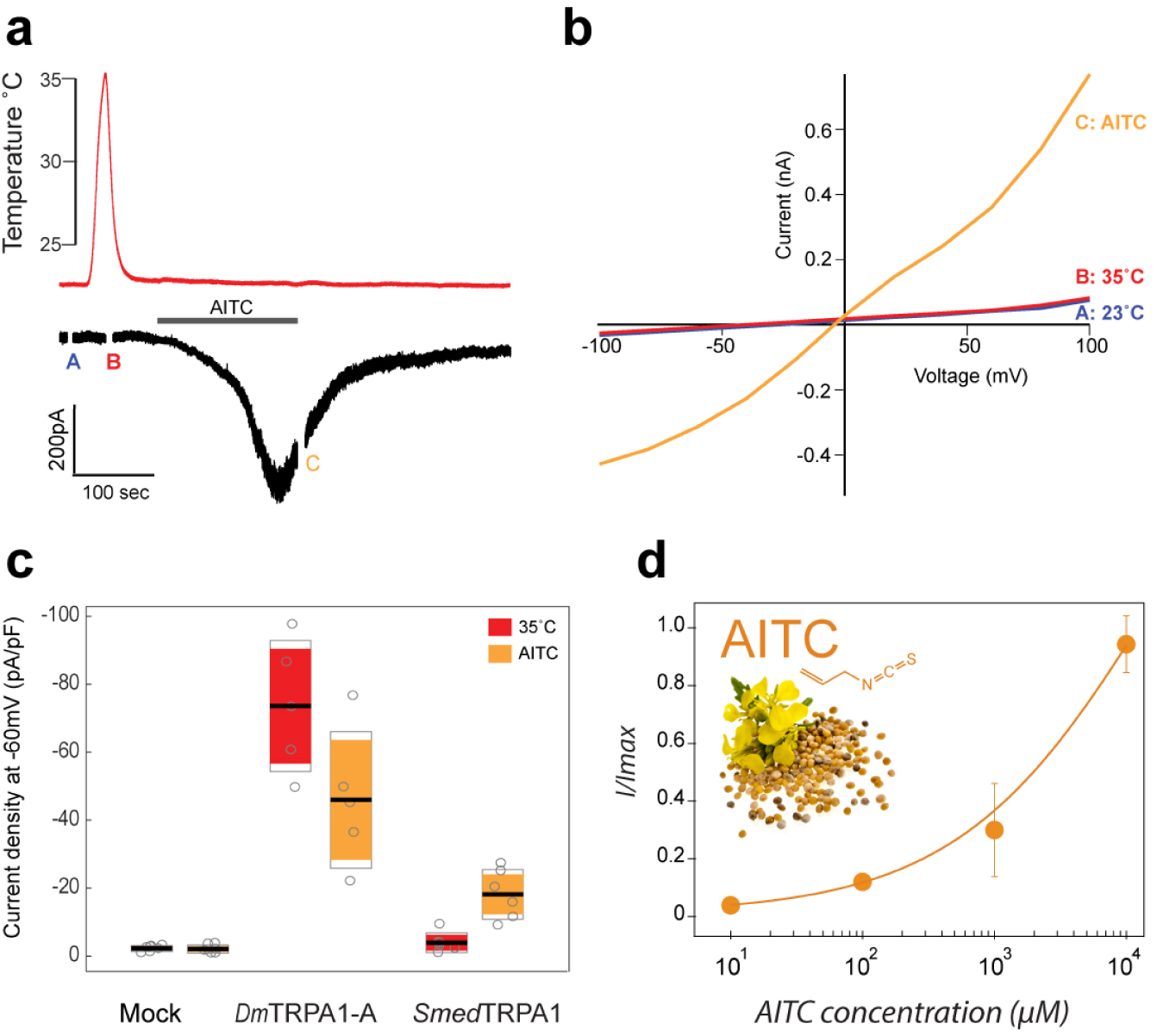
*Smed-TRPA1* expressed in *Drosophila* cells is activated by AITC but not by heat. **a**) S2R+ *Drosophila* cells voltage clamped at-60mV were stimulated by heat (red trace) and by bath application of AITC (500μM, grey bar). AITC application (but not heating) resulted in an inward current. **b**) Current/Voltage relationship from averages of three step protocols done at room temperature (blue trace), during the heat stimulation (red trace), and at the end of AITC application (orange trace; note that the timing of each set of measurements is also labeled as **A,B** and **C** on the trace shown in panel A). **c**) Current density (Max/capacitance) at 32°C and in the presence of AITC recorded in mock-transfected, *dTRPA1-A* transfected, and *Smed-TRPA1* transfected cells. Black line = mean; Colored boxes = +-STD; Grey empty boxes = 95% Confidence Interval. **d**). Dose-response for AITC activation of Smed TRPA1 (AV±STD; n=5 cells/condition; mustard flower and seed represent an iconic source of AITC).

**Figure 4:**
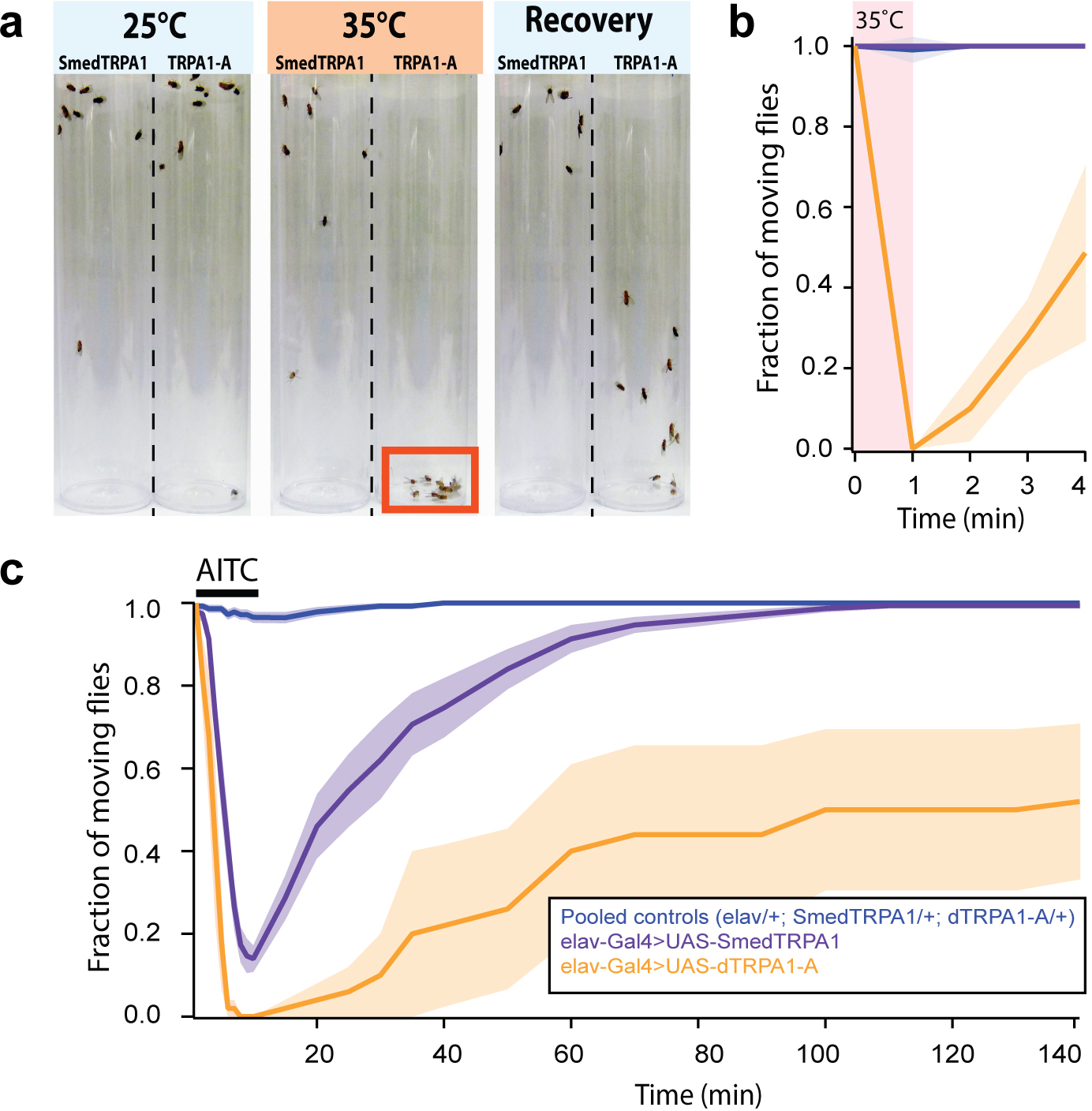
Functional expression of Smed-TRPA1 *in vivo* in adult *Drosophila* further demonstrates that the channel is sensitive to AITC but not to heat (°C). **a**) Adult fruit flies expressing either Smed-TRPA1 or–as a control-the intrinsically heat sensitive *Drosophila* TRPA1-A splice variant, throughout the nervous system (under the control of elav-Gal4) were subjected to a brief step at 35°C (a temperature which does not normally impair fly activity). TRPA1-A expressing flies are readily and reversibly incapacitated by heat (presumably because of simultaneous depolarization of neurons, caused by channel opening) and fall to the bottom of the tube; Smed-TRPA1 flies appear instead unaffected. **b**) Quantification of the experiment in **a**. Blue trace = pooled controls (*elav/+*; *UAS-Smed-TRPA1/+*; *UAS-TRPA1-A/+*; N=4 groups of 10 animals for each, tested separately); purple trace = experimental animals (*elav-Gal4>UAS-Smed-TRPA1*; N=4 groups of 10 animals); orange trace = positive control (*elav-Gal4>UAS-TRPA1-A*; N=4 groups of 10 animals; for all traces shaded area ±SEM). **c**) Adult flies expressing either Smed-TRPA1 or TRPA1-A were reversibly incapacitated by brief exposure to AITC vapors (see methods for details; Groups and Ns as above; shaded area ±SEM).

The lack of thermal sensitivity of *Smed-TRPA1 in vitro* (vis-à-vis the effect of RNAi on noxious heat avoidance) appears puzzling. However, TRPA1 is well known to function both as a primary temperature receptor as well as a signal transduction component, i.e. *downstream* of diverse signaling events ^10,33^. Interestingly, both the heat-nociception phenotype of TRPA1 mutant fly larvae and heat-entrainment defects of adults were readily rescued by a non-heat-sensitive variant of the fly TRPA1 (TRPA1-C, ^3,34^). These observations led us to test directly the possibility that the non-heat-sensitive Smed-TRPA1 may also be able to substitute for the fly TRPA1–i.e. to attempt across-phylum rescue of a *Drosophila* TRPA1 mutant by expression of the planarian homolog.

First, we used a rapid 2-choice assay for temperature preference ^28^ (similar to that described above), and probed the responses of wild type and TRPA1 *Drosophila* mutants to both innocuous (30°C) and noxious heat (40°C, Figure 5a). Consistent with previous reports ^2^, in our assay TRPA1 mutant flies showed a clear defect in the avoidance of noxious heat (Figure 5a,c; note that residual heat avoidance is likely mediated by *GR28b.d*-a distinct molecular hot receptor ^35^). Much like in the larva, this nociceptive phenotype could be significantly rescued by ubiquitous expression of *TRPA1-C*, a TRPA1 variant not directly activated by heat ^3^ (Figure 5b,c). Strikingly, a comparable amount of rescue could be achieved by ubiquitous expression of the planarian *Smed-TRPA1* (34% identical and 53% similar to the fly TRPA1 in amino-acidic sequence; Figure 5b,c), and even of a human *TRPA1* cDNA (36% and 31% identical, and 57% and 49% similar to fly and Smed-TRPA1, respectively; Figure 5b,c). The rescue of the fly phenotype by an evolutionary distant ‘heat-insensitive’ homolog such as *Smed-TRPA1*, and the human TRPA1 (activated by cold rather than heat), argues that the function of TRPA1 in heat nociception is unlikely to be fully explained by direct heat gating.

**Figure 5:**
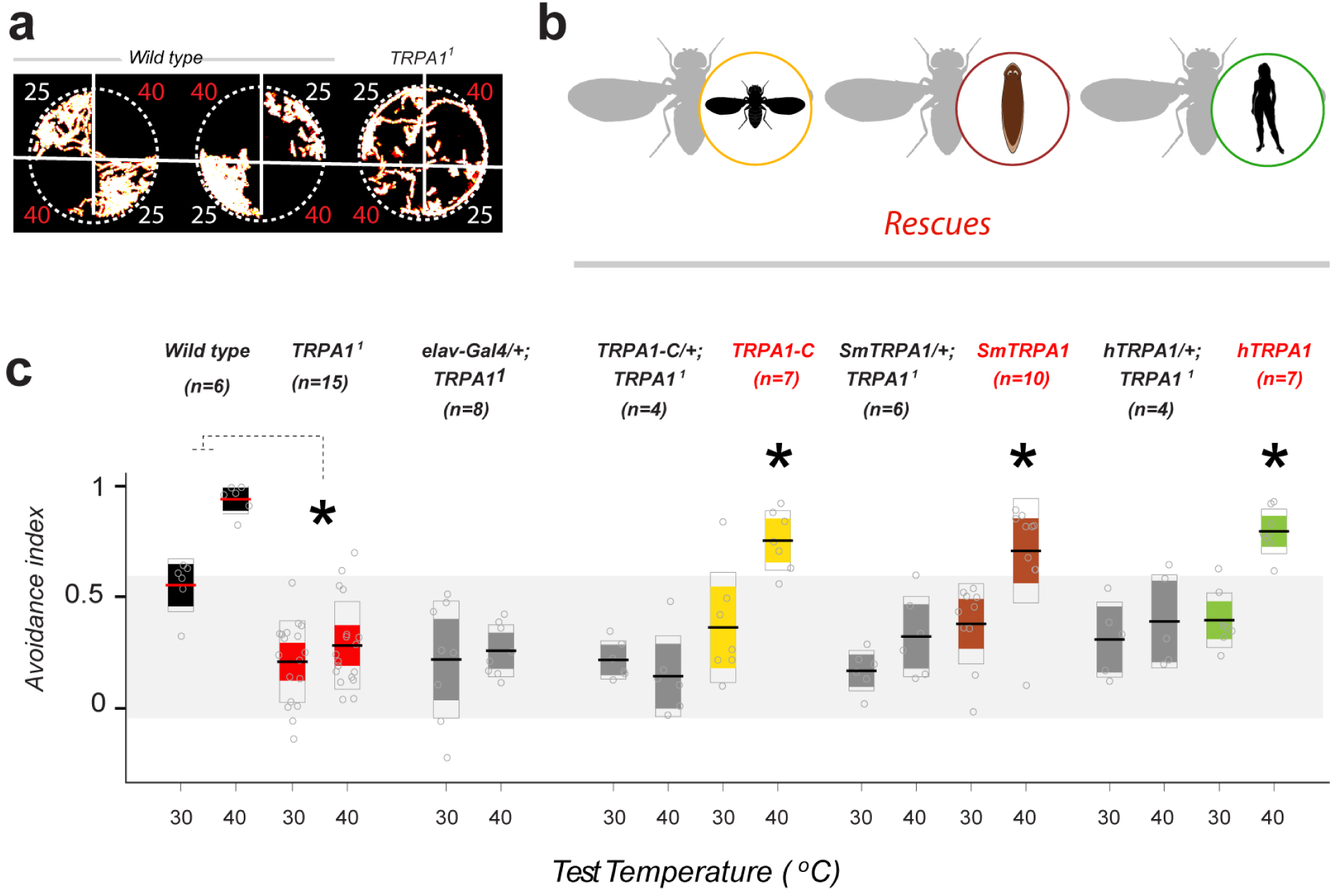
Across-phylum rescue of *Drosophila* TRPA1 mutant phenotypes by planarian and human TRPA1. **a**) In a 2-choice assay, wild type *Drosophila* flies robustly avoid noxious heat (40°C). In contrast, TRPA1^1^ mutants explore more readily the 40°C quadrants (the panels are maximum-projections of 3′ movies, illustrating the extent of fly movement, temperature in °C is indicated next to each quadrant). **b**) Schematic of the rescue experiments. **c**) Avoidance index of wild-type (black boxes), TRPA1^1^ mutants (red), rescues (yellow, brown, green), and control genotypes (grey). TRPA1^1^ mutants display a significantly lower avoidance index for heat. Pan-neural expression (under the control of elav-Gal4) of mRNA encoding *Drosophila* TRPA1-C (a splice variant encoding a channel that is not heat-sensitive, yellow), Smed-TRPA1 (brown), or human TRPA1 (green, each under a UAS-promoter) significantly rescues noxious heat avoidance (40°C). Control genotypes: elav driver/+; TRPA1^1^ and UAS-transgene/+; TRPA1^1^ (see methods for full genotypes). Avoidance index values for each test temperature were compared by unpaired t-tests (wt vs TRPA1^1^, P<0.001) or two-way ANOVAs (for all other genotypes), were asterisks denote a significant interaction between the Gal4 and UAS transgene (* = P<0.01). Thick line = mean; Colored boxes = +-STD; Grey empty boxes = 95% Confidence Interval.

Instead, a number of observations point towards early markers of tissue damage as potential mediators of TRPA1 activation during nociceptive heat responses. Hydrogen peroxide (H_2_O_2_), is amongst the earliest known markers of mechanical tissue damage in vertebrates ^36^, *Drosophila* ^37^ as well as *Planarians* ^38^. H_2_O_2_ is a well-known activator of mammalian and *Drosophila* TRPA1 (^39-43^, together with additional Reactive Oxygen Species–ROS ^44^), and recent work suggest that responses to potentially damaging short-wavelength UV light occurs through photochemical production of H_2_O_2_, and requires TRPA1 in both flies ^41-43,45^ and Planarians ^46^. Thus, if noxious heat were to cause rapid, localized, production of H_2_O_2_/ROS, this could provide the direct signal for TRPA1 activation that mediates nociceptive responses.

For this hypothesis to be correct, a number of conditions have to be met: (1) *SmedTRPA1* (like human and *Drosophila*) should be activated by H_2_O_2_; (2) *In vivo*, heat stimulation in the appropriate range should cause rapid H_2_O_2_/ROS production (on a timescale compatible with the animal’s escape behaviors) and, (3) if indeed nociceptive heat responses are mediated by H_2_O_2_/ROS, an acute increase in H_2_O_2_/ROS levels should sensitize the animal’s behavioral responses to noxious heat, and this sensitization should depend on TRPA1. Our experiments confirm each of these predictions.

First, we tested Smed-TRPA1 for potential responses to H_2_O_2_ *in vitro* in our cell expression system (see above). Our recordings showed that, in S2 cells, Smed-TRPA1 was indeed activated by a range of H_2_O_2_ concentrations (Figure 6a,b), as was the *Drosophila* counterpart TRPA1-C (Figure 6b; i.e. the ‘heat insensitive’ fly variant which also supported behavioral rescue in our experiment). We note that, while it is difficult to speculate on the H_2_O_2_/ROS concentration that TRPA1 may encounter during a heat challenge *in vivo*, secondary modifications (such as prolyl hydroxylation ^47^) have been shown to dramatically increase TRPA1 responses to H_2_O_2_/ROS, potentially expanding the effective sensitivity of the channel.

**Figure 6:**
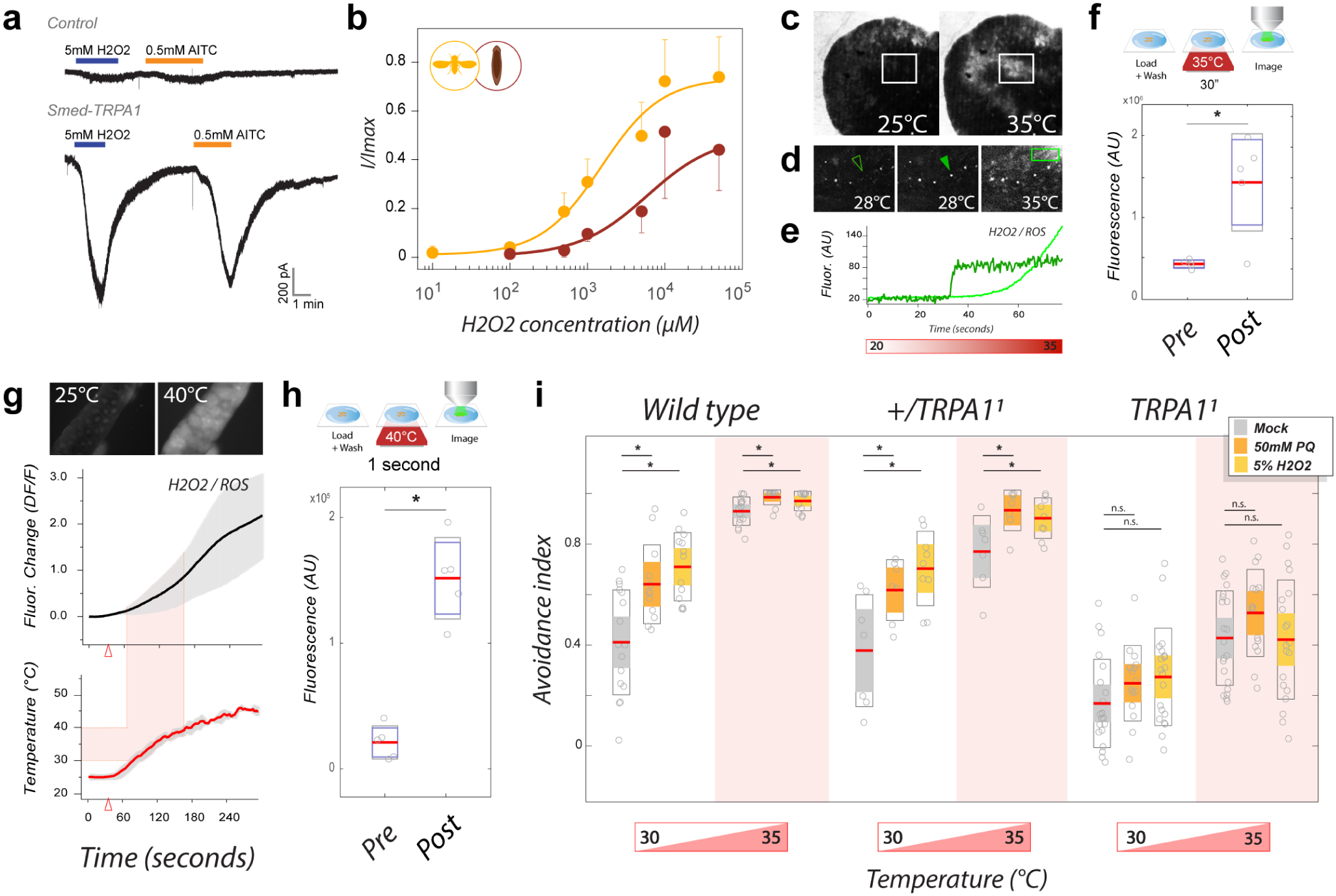
H_2_O_2_/ROS as a signal for TRPA1 activation during noxious heat responses. **a**) Heterologously expressed *Smed-TRPA1* is activated by H_2_O_2_. **b**) Dose-response of H_2_O_2_ activation for *Drosophila* TRPA1-C (yellow trace and points), and Smed-TRPA1 (brown trace and points, AV±STD, n=5 cells/condition). **c-e**) The ROS dye Carboxy-H_2_DCFDA demonstrates in vivo ROS production in response to heat in living planarians. **c** and **d**) Representative frames of tissues and cells undergoing rapid fluorescent changes in response to heat. **e**) An ROI around the cell in **d** (arrow) and the square, light green box are plotted as traces. **f**) Exposure of planarian worms to 35°C for 30” results in a significant increase in fluorescence. **g**) Carboxy-H_2_DCFDA fluorescence in response to heating in *Drosophila* salivary gland tissue (AV±SD). **h**) Exposure of salivary glands to 40°C for 1” results in a significant fluorescence increase (in **f** and **h**, * = P<0.05; n=5/condition). **i**) Acute feeding with pro-oxidants sensitizes adult *Drosophila* to heat. Paraquat (orange bars) or H_2_O_2_ (yellow bars) feeding, results in increased heat voidance in a 2-choice behavioral assay in both wild-type and heterozygous TRPA1/+ controls (* = P<0.05). In contrast, heat avoidance does not increase in TRPA1 mutants (n.s. =not significant; n = 11-21. In **f**, **h** and **i**: Red line = mean; Colored boxes = +-STD; Grey empty boxes = 95% Confidence Interval).

Next, we tested if heat stimulation in the appropriate range (i.e. in the ‘noxious’ range for each species) may lead to the rapid production of H_2_O_2_/ROS in both *Schmidtea* and *Drosophila* tissue. For this experiment, we loaded living samples with the dye 5-(and 6)-carboxy-2’,7’-dichlorodihydrofluorescein diacetate (carboxy-H_2_DCFDA, a widely used fluorogenic ROS marker for live cells ^48^), and monitored potential fluorescence changes in response to heat exposure by confocal microscopy. As previously reported, live planarian worms could be directly loaded with the dye, and displayed H_2_O_2_/ROS induced fluorescence at sites of physical wounding (^38^ and data not shown; and see **ED** Figure 3 for additional controls). Fast H_2_O_2_/ROS production was also detected when the worms were submitted to rapid heating while under the microscope (i.e. by using a heating stage; Δt=20>35°C, speed=∼1°C/4 seconds). Consistent with the ‘noxious’ range for this animal (30-35°C), we observed rapid H_2_O_2_/ROS increases starting above 23-25°C, and culminating in widespread fluorescence around 30-35°C (Figure 6c-e). *Drosophila* tissues also displayed rapid H_2_O_2_/ROS increases in response to heating, but this time the increase in fluorescence started around 30°C, and culminated around 40-45°C (Figure 6f-g), consistent with the ‘noxious’ range for *Drosophila* (35-40°C). Importantly, in both planarians and fly tissues, we recorded fluorescence changes rapid enough to be compatible with the timescale that would be required to trigger/modulate behavioral responses: for example between two imaging frames (Figure 6d, i.e. separated by ∼300ms), or after as little as one second of exposure to heat (Figure 6h).

Carboxy-H_2_DCFDA is a general oxidative stress indicator, and does not discriminate amongst different reactive oxygen species. To directly test if H_2_O_2_ in particular may be amongst the species produced during a noxious heat challenge, we turned to the genetically encoded H_2_O_2_ indicator roGFP2-Orp1. This indicator couples the redox-sensitive green fluorescent protein 2 (roGFP2) with the yeast H_2_O_2_ sensor Orp1 ^49^, allowing the measurement of changes in H_2_O_2_ levels in intact living animals ^50^. Our results show that, in transgenic *Drosophila* larvae, roGFP2-Orp1 reported a significant increase in H_2_O_2_ upon brief (∼5 seconds) exposure to noxious heat (Figure ED4, see legend and methods for details).

Finally, we tested the notion that–if noxious temperatures are indeed sensed at least in part through H_2_O_2_/ROS production-an acute, systemic increase in H_2_O_2_/ROS levels may sensitize the animal’s behavioral responses to heat. Here, we tested adult *Drosophila* for heat avoidance using our 2-choice assay (see above), but this time the flies’ performance in the arena was preceded by a short feeding with H_2_O_2_ ^51^ or Paraquat (a potent pro-oxidant ^52^). Strikingly, pro-oxidant feeding significantly increased heat avoidance scores in both wild type and control flies (to both 30° and 35°C) but not in TRPA1 mutant flies (Figure 6i). This result directly demonstrates that H_2_O_2_/ROS can sensitize aversive responses to heat in *Drosophila*, and that this sensitization requires functional TRPA1 channels.

Planarian flatworms are a powerful, and yet underutilized model for behavioral research. They are capable of active hunting behavior, and possess simple sensory systems ^30,53^ and a simple brain that operates using synaptic and neurotransmitter principles similar to those of the more complex insect or mammalian brains ^54^.

Here, we have shown that the ion channel TRPA1 functions as a key transduction component for nociceptive signals in *Schmidtea mediterranea*. In insects and many vertebrates (snakes, frogs etc.), TRPA1 channels can be directly gated by temperature changes, but our work in *Schmidtea* and *Drosophila* suggests that TRPA1’s function in nociceptive heat sensing goes beyond that of a canonical heat-activated ion channel. Instead, our data suggest that H_2_O_2_/ROS are rapidly produced in response to noxious heat, and that this signal contributes to channel activation to mediate defensive responses. We note that this mechanism provides an especially satisfactory explanation for the remarkable across-phylum rescue of heat-avoidance phenotypes of TRPA1 mutant *Drosophila* by both *Schmidtea* TRPA1 (insensitive to heat), as well as human TRPA1 (activated by cold). The heat range that is expected to produce tissue damage in *Drosophila* (i.e. the “noxious range”, ∼35-40°C) is different from that of *Schmidtea* (∼30°C) from that of human (∼45°C)-and yet transgenic rescue of fly mutants restored heat avoidance to that of the host (*Drosophila*) rather than the channel donor (*Schmidtea* or human). This can be easily accounted for by the fact that the thermal range that causes heat damage (and therefore H_2_O_2_/ROS production) is determined by the heat tolerance of the host tissue.

Finally, our results suggest that early Bilaterians already possessed a polymodal (chemical/thermal) nociceptive system that relied on H_2_O_2_/ROS-mediated TRPA1 activation. This core function has been conserved in extant lineages, and may have placed TRPA1 in a key position to undergo the additional transitions into a hot-or cold-activated channel that have been documented in different animal groups. Our results also imply that human pain systems may share a common ancestry with the nociceptive systems of extant bilateral animals, tracing back their origin to the common ‘Urbilaterian’ progenitor that lived more than 500 million years ago.

## Methods

### Cloning of a Smed-TRPA1 full length coding sequence

A full-length *Smed-TRPA1* coding sequence was amplified by PCR starting from a *Schmidtea mediterranea* cDNA library. The library was generated by Superscript III reverse transcription (Life Technologies) from total RNA, purified from whole animals using Trizol followed by DNAse treatment with DNA-free (Ambion/ThermoFisher). The following primers were used for PCR: FWD 5′-CAaaac<u>ATG</u>AATAAAATTTCTAAAAACCGAAAAACCTC-3’ and REV 5′-TTAAAAATTGTTATCTGGTTTGACAGATTTCTG-3’ (Kozak sequence for Drosophila in lower case letters). Analysis of 5′-and 3′-RACE libraries produced with SMARTer® RACE 5′/3’ Kit (Clontech) confirmed that the identified sequence includes the appropriate ATG and stop codons and represents a single, full-length Smed-TRPA1 coding sequence (3510 bp). Analysis of available RNAseq data (SmedGD, http://smedgd.neuro.utah.edu/; see Robb et al., Genesis. 2015 Aug;53(8):535-46) indicates that *Smed-TRPA1* likely produces a single transcript, encoding a protein of 1169 amino acids, which contains 14 N-terminal ankyrin repeats (Interpro database entry: IPR002110) followed by an ion transporter domain (Interpro database entry: IPR005821).

### Phylogenetic analysis of TRPA1 homologs

A neighbor-joining phylogenetic tree was constructed using the full sequence of *bona fide* (experimentally validated) TRPA1 proteins from 25 organisms: *A. gambiae* (ACC86138), *A. aegypti* (AAEL009419), *D. melanogaster (AE*U 17952), *B. mori (NP_00*1296525), *C. elegans* (ABQ15208), *C. brevicauda* (AEL30802), *C. porcellus* (NP_001185699), *C. hortulanus* (ADD82932), *C. atrox* (ADD82930), *D. rerio* (NP_001007066 and NP_001007067), *D. rotundus* (AEL30803), *G. gallus* (NP_001305389), *H. armigera* (AHV83756), *H. sapiens* (NP_015628), *M. mulatta* (XP_001083172), *M. musculus* (NP_808449), *P. obsoletus lindheimeri* (ADD82929), *P. jerdonii* (AEW26660), *P. regius* (ADD82928), *R. norvegicus* (NP_997491), *T. rubripes* (XP_003968031), *T. castaneum* (LOC658860), *V. destructor* (BAO73033 and BAO73034), *X. tropicalis* (BAM42680).

### Planarian RNAi

The CIW4 asexual laboratory strain of *Schmidtea mediterranea* was used for all experiments. Animals were kept in plastic containers filled with 1x artificial planarian water (APW) that contained: 1.6mM NaCl, 1mM CaCl_2_, 1mM MgSO_4_, 0.1mM MgCl_2_, 0.1mM KCl, 1.2mM NaHCO_3_. Planarians were fed with homogenized calf liver for stock maintenance. The containers were cleaned two days after feeding or once a week if starved. Templates for RNA synthesis were generated by PCR from pGEM-t vectors (Promega) harboring 1.5kb fragments of Smed-TRPA1 or Smed-AP2 cDNA; a 0.8kb UNC22 PCR product was used to generate UNC22 template as an RNAi control (UNC22 is a *C. elegans* gene not present in the *S. mediterranea* genome). The T7 RNA polymerase promoter was introduced to the 5′end or 3′end of the corresponding fragment, and 2 subsequent PCR reactions were performed to generate sense and antisense RNA strands. Sense and antisense strands were pooled together, purified by phenol-chloroform extraction, and resuspended in 16μl of water before being annealed by incubating at 72°C then 37°C and finally on ice. dsRNA was mixed with 80μl of homogenized calf liver and 2μl of red food coloring to assess food intake. Planarians were starved for at least a week and then fed the dsRNA every other day 3-4 times (15μl of food for 10-15 animals). Animals that did not feed were discarded. For behavioral experiments animals were used between one day and four days after last feeding, phototaxis was used at the end of each experiment to ensure viability.

### Expression analysis by Q-PCR

Total RNA from UNC22, Smed-AP2 and Smed-TRPA1 knockdown planarians was purified using Trizol reagent (Life Technologies). First strand cDNA was synthesized using MultiScribe Reverse Transcriptase (Fisher Scientific) from DNAse-treated (TURBO DNAse, Ambion) total RNA. qPCR reactions were performed using the EvaGreen dye (Biotium). 4 biological replicates were run for each treatment, with clathrin mRNA detected as reference gene for quantifying expression changes using the delta-delta Ct method and normalizing to the expression obtained in the control RNAi treatment. *Smed-TRPA1* was detected using the primers 5′-ACTCTCATCAACAGACAGACTTGT-3’ and 5′-ATTTCAGCCTCTGGATCCATTTCC-3’ and *clathrin* primers were 5′-GACTGCGGGCTTCTATTGAG-3’ and 5′-GCGGCAATTCTTCTGAACTC-3’. Results were compared using Kruskal-Wallis one-way ANOVA.

### Fluorescent *in situ* hybridization (FISH)

*Smed-TRPA1* riboprobes were generated from a PCR fragment flanked by T7 promoter sequences using RNA DIG-labeling mix (Sigma-Aldrich). After *in vitro* transcription, antisense probes were precipitated with 100% ethanol and resuspended in 25μl of deionized formamide. Planarians were killed in 5% N-Acetyl cysteine, fixed in 4% formaldehyde, followed by dehydration and overnight bleaching in 6% H_2_O_2_ on a light box. Animals were preserved in 100% methanol and stored at-20°C. For FISH, planarians were re-hydrated with a methanol:PBST (PBS, 0.1% triton X-100) dilution series; next, animals were treated with 10mg/ml proteinase K, post-fixed in 4% formaldehyde, and incubated at 56°C for 2 hours in pre-hybridization solution (50% of de-ionized formamide, 5x SSC, 0.1 mg/ml yeast RNA, 1% Tween-20 in DEPC-treated water). Hybridization with riboprobes was conducted for 16h in hybridization solution (same as pre-hybridization solution plus 5% dextran sulfate). Then animals were washed in pre-hybridization solution, and then subjected to a dilution series of 2X SSC, then 0.2X SSC, and finally TNTx (0.1 M Tris pH 7.5, 0.15 M NaCl, 0.3% Triton X-100). Animals were blocked in TNTx plus 5% horse serum and 5% Western Blocking Reagent (Sigma-Aldrich) for 2h at RT, and then labeled with a sheep anti-DIG-POD antibody (1:2000, Sigma-Aldrich) in blocking solution overnight at 4°C. Animals were washed 8X in TNTx, incubated in Tyramide solution with rhodamine (1:500) and H_2_O_2_ for 10 minutes with shaking. Finally, animals were rinsed 6X in TNTx. ISH experiments were performed four times with similar results. To quantify TRPA1+ cells in various groups (Figure 1F-K), ten worms per treatment were imaged with a Leica DM 2500 confocal microscope with a 10X objective and 1.5 digital zoom, and Z-stacks encompassing the thickness of each animal were acquired at 5 μm intervals, using constant laser and PMT settings. Z-stacks were analyzed using ImageJ: Brightness/Contrast was adjusted in batch using identical settings and max projection images through the animal were produced. From these, the number of fluorescent cells in a defined ROI in the brain region and a defined ROI at the periphery (each of constant size, shown as yellow boxes in Figure 1) were counted. The number of fluorescent cells was then plotted and compared by unpaired t-test.

### Planarian behavioral assays: heat avoidance

Heat avoidance was measured in the “Planariometer” (see Supplementary Figure 1). The Planariometer consisted in four independent tiles covered by thin anodized aluminum foil. A hydrophobic ink pen (Super PAP pen – ThermoFisher) was used to create a circular barrier (55mm of diameter) to allow a thin film of water (1-2 mm) to form a central pool in which the worms can move freely. In each experiment, 2 opposite tiles were set at 32°C and 2 at 24°C and animal movement was recorded for 4 minutes. The spatial configuration of hot and cool tiles was then reversed for 4 additional minutes (and a second movie acquired) to control for potential spatial biases. Experiments were conducted in the dark with infrared (IR) LED illumination and videos were recorded with an IR-sensitive CCD camera (Basler). Five independent groups of 10 animals per treatment were used. The heat avoidance index (AI = #worms at 24°C-#worms at 32°C/total #worms) was calculated from the last 120 frames of each video (last minute of the video) by measuring the positions of the worms every 3 frames using a custom-made Matlab script. Avoidance index values were compared using Kruskal–Wallis ANOVA followed by Tukey’s honest significant difference test.

### Planarian behavioral assays: AITC avoidance

Five independent groups of 10 animals per treatment were tested in an arena composed of two circular chambers connected by a narrow corridor. At the beginning of the experiment, all animals were placed in chamber 1 together with a small block of control agar (1% agarose dissolved in 1x APW) or AITC laced agar. The AITC laced agar was made with 1% agarose in 1x APW and 50mM AITC. The chamber was placed in the dark, animals where illuminated with IR light and their behavior was recorded for 5 minutes with a IR-sensitive CCD camera. Videos were analyzed and the number of planarians in each chamber was quantified every 10 frames for the last 125 frames of the video (last minute). The fraction of animals in each chamber was counted after each treatment and compared using Kruskal–Wallis ANOVA followed by Tukey’s honest significant difference test.

### Planarian behavioral assays: Negative phototaxis

Four independent groups of 10 worms per treatment were placed in chamber 1 of the arena as described above (AITC avoidance experiments). The arena was either kept completely dark (control condition), alternatively, chamber 1 was exposed to bright light while chamber 2 was kept dark. Animals were allowed to distribute for 2 minutes before the number of worms in each chamber was counted.

### Cell transfections

pAC-GFP, pAC-Smed-TRPA1 and pAC-dTRPA1-A were generated by cloning GFP, *Smed-TRPA1* ORF (see above) and a dTRPA1-A cDNA (a gift of Dan Tracey) into pCR^™^8/GW/TOPO® TA (ThermoFisher) and then transferring them into pAC-GW expression vector [created by ligating the Gateway cassette from pMartini Gate C R2-R1 (Addgene plasmid #36436) cut with XhoI and XbaI into pAc5.1/V5-His A (ThermoFisher)]. S2R+ cells (a gift from R. Carthew) were cultured in Schneider’s Drosophila Medium (Lonza) supplemented with 10% fetal bovine serum and 1% penicillin-Streptomycin mixture (100units/mL and 100μg/mL respectively, Fisher Scientific). For electrophysiological recordings, S2R+ cells where grown on coverslips in Schneider’s Drosophila Medium supplemented with 50μM LaCl_3_ and transfected with 50ng of pAC-GFP vector and 500ng of either pAC-dTRPA1-A or pAC-Smed-TRPA1 vectors mixed with 4μl of enhancer and 150μl of buffer EC. After 5 min, 6.5μl of Effectene® Transfection Reagent (Qiagen) was added and the mix was incubated for 10min before being dispensed to the cells. Transfected cells were incubated at RT for at least 36h to allow gene expression.

### Electrophysiological recordings

Whole cell voltage clamp was performed on S2R+ transfected cells identified by GFP fluorescence. The intracellular solution contained: 140mM methanesulfonic acid, 2mM MgCl_2_, 1mM EGTA, 5 mM HEPES, 1 mM Na2ATP; pH was adjusted to 7.3 with CsOH and the osmolarity was adjusted to 315 ± 5mOsml with sucrose. The extracellular solution contained: 140mM NaCl, 5mM KCl, 1mM CaCl_2_, 1mM HEPES, 10mM Glucose; pH was adjusted to 7.2 with NaOH and the osmolarity was adjusted to 310 ± 5 mOsml with sucrose. Patch pipettes resistance ranged from 5 to 10MΩ. Recordings were obtained with an AxoPatch 200b amplifier (Axon Instruments), and analyzed with AxoGraph software and custom-made Matlab scripts. Recordings were made with 1x output gain and 5KHz low pass filter. Bath offset and capacitance were compensated; series resistance was 9.5±5.5MΩ without compensation. Recordings were made at RT (22-23°C) and temperature stimulation was achieved by raising the temperature of the bath solution via an inline heater (HPT-2A, ALA Scientific Instruments) and a TC-20 temperature controller (NPI Electronics). Temperature was monitored with a T-384 thermocouple (Physitemp Instruments) tethered to the electrode holder, so that the tip of the thermocouple was approximately at a constant distance from the tip of the recording electrode (1-2 mm). Chemical stimulation was achieved by bath perfusion of extracellular solution containing 500μM of allyl isothiocyanate (AITC, Sigma). Cells were held at-60mV and currents were monitored during heat and chemical stimulation. Current-voltage relationships were constructed by averaging three step protocols consisting of 100ms steps of 20mV from-100 to 100mV separated by 400ms. These IV relationships where made at RT, during the heat stimulation, and at the end of a 3min AITC application. Note that Smed-TRPA1 did not appear to respond to cooling. For the AITC and Hydrogen peroxide dose responses (H_2_O_2_, Sigma, 30% w/w) we used 1 min stimulation at each concentration. Recordings were performed as described above with the exception that the intracellular solution contained 140mM K-gluconate instead of Cesium.

### Fly strains and transgenes

Flies were reared on standard cornmeal agar medium at room temperature (RT). The following fly strains were used: Canton-special, isogenic w^1118^ (a gift from Marcus C. Stensmyr); elav-Gal4/CyO; *TRPA1^1^* (BDSC#26504, backcrossed 5 times); 5xUAS-TRPA1-C (a gift from D. Tracey); tub-cyto-roGFP2-Orp1 (ref #50). To generate *UAS-SmedTRPA1* flies, *Smed-TRPA1* cDNA (see above) was cloned into pCR^™^8/GW/TOPO® TA (Invitrogen) and then transferred into a 40xUAS destination vector created introducing the Gateway® cassette into pJFRC8-40XUAS-IVS-mCD8::GFP (Addgene #26221) via the XhoI/ XbaI restriction sites. This construct was then used for embryo injection by BestGene Inc. to generate P[40XUAS::Smed-TRPA1]attP40 flies. Similarly, *UAS-humanTRPA1* flies were obtained starting from a human TRPA1 cDNA (NP_015628; a gift from Mark Hoon) to generate P[40XUAS::hTRPA1]attP40 flies. Expression of the transgenes was confirmed by RT PCR. Full genotypes of fly stocks used in Figure 5: *w^1118^. w^1118^; TRPA1^1^. w^1118^; elav-Gal4/+; TRPA1^1^/TRPA1^1^. w^1118^; +/UAS-TRPA1-C; TRPA1^1^/TRPA1^1^. w^1118^; elav-Gal4/UAS-TRPA1-C; TRPA1^1^/TRPA1^1^. w^1118^; +/UAS-Smed-TRPA1; TRPA1^1^/TRPA1^1^. w^1118^; elav-Gal4/UAS-Smed-TRPA1; TRPA1^1^/TRPA1^1^. w^1118^; +/UAS-human TRPA1; TRPA1^1^/TRPA1^1^. w^1118^; elav-Gal4/UAS-human TRPA1; TRPA1^1^/TRPA1^1^*.

### *Drosophila* behavioral assays

Temperature preference assay were performed as previously described (Gallio et al., 2011). In brief, Avoidance Index (AI) values for the test temperatures (TT) are calculated as follows: AI=# flies at 25°C-# flies at TT/(total # flies). AI values were compared using ANOVA or 2-way ANOVA as previously described. Avoidance index values for experiments with Gal4 and UAS lines were compared by two-way ANOVA (threshold P = 0.01). Kolmogorov Smirnov tests were used to test for a normally distributed sample. Homogeneity of variance for each data set was confirmed by calculating the Spearman correlation between the absolute values of the residual errors and the observed values of the dependent variable (threshold P = 0.05). Statistical analysis was carried out in MATLAB; sample sizes were chosen to reliably measure experimental parameters. Experiments did not involve randomization or blinding. All temperature preference experiments were performed in a custom chamber kept at a constant RH of 40%. Heat and AITC-vapor incapacitation experiments (Figure 4) were performed as follows: Flies of the genotypes elav/+, UAS-Smed-TRPA1/+, UAS-TRPA1-1/+ (negative controls), elav-Gal4>UAS-TRPA1-A (positive control) and elav-Gal4>UAS-Smed-TRPA1(experimental animals), were used to test responses to temperature and AITC. For heat incapacitation, groups of 10 flies were collected in empty vials and placed in a 25°C incubator for at least an hour before the experiments. Next, the vials were submerged in water (pre-heated to 35°C) and kept submerged until the internal air temperature of the tube had been at 35°C for one minute (as measured by a thermocouple). Following this, the number of incapacitated flies (i.e. flies that had dropped to the bottom of the tube) was counted. Vials were then placed at RT for three additional minutes, and the number of incapacitated flies scored every minute to measure recovery. For the exposure to AITC vapors, groups of 10 flies of each genotype (see above), were collected in 15mL tubes for bacterial culture with small holes to allow air flow. These 15mL culture tube were placed inside a 50mL conical tube containing a small piece of filter paper with 1μl of 2.5M AITC. Flies were exposed to AITC vapors for 10minutes and then transferred to clean vials for recovery. The number of incapacitated animals was recorded every minute during AITC exposure and every 5 minutes during recovery. For the pro-oxidant feeding experiments, groups of twenty 3-5 day-old flies of the appropriate genotype were starved for 18 hours in vials with a Kim-wipe saturated by water. Flies were then fed for three hours on Nutri-Fly^™^ Instant Medium (Genesee Scientific #66-117) prepared with the respective pro-oxidant solution at a ratio of medium to liquid of 1:3. The liquid used contained the pro-oxidant and sucrose, or sucrose alone (’mock’). Final concentrations: all samples=2% sucrose; H_2_O_2_=5%, Paraquat (Sigma #856177)=50mM. Immediately after, the animals were tested for temperature preference as described above. Food intake was monitored in parallel experiments using green food colorant (25μl for 3ml of solution).

### ROS imaging

To evaluate ROS levels in response to heat stimuli in intact live planarians and *Drosophila* tissues, we used the fluorogenic oxidative stress indicator carboxy-H2DCFDA (Molecular Probes #I36007) per the manufacturer’s guidelines (and see below). ROS levels were imaged on an LSM510 Zeiss confocal microscope equipped using a 488 argon laser. Temperature stimuli were generated with a Model 5000 KT stage controller (20/20 Technology) and the temperature was recorded using NI USB-TC01 equipped with a thermocouple probe (National Instruments). Intact planarians were incubated for an hour in 25μM carboxy-H2DCFDA (diluted in APW) and washed briefly in APW prior to imaging. To minimize movements during scanning, animals were placed into a tight-fitting custom made Sylgard frame mounted on a glass slide and filled with APW. The preparation was sealed with a cover slip. For real time ROS detection, planarians were imaged continuously with a 5x/0.16 Zeiss air objective at 256x256 pixel resolution and 2x optical zoom at 0.395msec frame rate during a heat ramp of Δt=20>35°C, speed=∼1°C/4 seconds. ROS induced fluorescence was measured from confocal images acquired at low resolution and using a fully open pinhole, from ROIs corresponding to large parts of the head region (512x512 pixel resolution 1x optical zoom with a 5xZeiss air objective). Drosophila salivary glands were dissected in PBS and incubated with 25μM carboxy-H2DCFDA (diluted in PBS) for an hour at room temperature prior to heat stimulation. Tissues were then briefly washed in PBS and transferred into a custom made thin Sylgard frame containing PBS and mounted on a glass slide. The preparation was sealed with a cover slip. For real time ROS detection in *Drosophila*, salivary glands were imaged continuously with a 10xZeiss air objective at 256x256 pixel resolution and 2x optical zoom at 0.395msec frame rate during a heat ramp of Δt=20>45°C, speed=∼1°C/4 seconds. DF/F analysis was carried out using custom scripts in MATLAB, base fluorescence was calculated using all frames preceding temperature trigger (occurring at 30s). Confocal stacks of about 100 μm at 5 μm steps were obtained at 512x512 pixel resolution 1x optical zoom with a 10x Zeiss air objective.

For in vivo H2O2 detection in intact animals, tub-cyto-roGFP2-Orp1 larvae were placed on a heated surface set to ∼35°C for 5 seconds. As a positive control, larvae were exposed to 25 μM H2O2 for 10 minutes. Animals were rapidly dissected post treatment in PBS to extract their wing imaginal discs (as described in ref #50). The tissues were mounted in glycerol and immediately imaged on a Leica SP5 inverted confocal microscope equipped with a 405 UV and a 488 Argon laser and a 10x air objective at 512x512 pixel resolution and 400 Hz. Image acquisition and processing were performed as above. We used two-sample T-test to calculate significant difference (p<0.05) between treatments and controls. Excitation of the biosensor fluorescence by the 405 nm and 488 nm laser lines was performed sequentially and stack by stack. Emission was detected at 500–570 nm. Image processing was performed with imageJ. Control fluorescence (i.e. of untreated tissue) was set to 1.

## Supplementary Information Line

Supplementary information is linked to the on-line version of the paper at www.nature.com

## Acknowledgments

We thank D. Tracey and R. Carthew for reagents. Andrew Kuang and Leah Vinson for technical assistance, Lindsey Macpherson, David Yarmolinsky and members of the Gallio Lab for comments on the manuscript, Indira Raman for technical advice and Marcus Stensmyr for the kind gift of the fly drawing in Fig1. Work in the Gallio lab is supported by NIH grant R01NS086859 (to M.G.), the Chicago Biomedical Consortium with support from the Searle Funds at the Chicago Community Trust (to E.E.Z.), and by training grant 2T32MH067564 (to O.M.A.). Work in the Petersen Lab is supported by an NIH Director’s New Innovator award (1DP2DE024365-01).

## Author contributions

M.G. designed the study, analyzed the data, and wrote the paper with critical input from all authors; O.M.A. performed all planarian behavioral experiments and electrophysiology and analyzed the corresponding data. E.E.Z. performed all fly rescue experiments and ROS assays and analyzed the corresponding data. A.P. cloned *Smed-TRPA1*, produced rescue constructs and transgenics and analyzed sequences with help from C.P.P; A.P. O.M.A and C.V.D. performed Q-PCR and ISH experiments. E.E.Z and O.M.A generated human-TRPA1 expressing flies.

## Author information

Reprints and permissions information is available at www.nature.com/reprints. The authors declare no competing financial interests. Correspondence and requests for materials should be addressed to M.G.

